# The aryl hydrocarbon receptor interacting protein suppresses RNA virus-induced type I IFN and IL-6 in mouse macrophages

**DOI:** 10.1101/2025.06.30.662398

**Authors:** Sarah A. Kazzaz, Todd M. Umstead, Zissis C. Chroneos, Edward W. Harhaj

## Abstract

Aryl hydrocarbon receptor (AhR) interacting protein (AIP) suppresses type I IFN production by interacting with and preventing the nuclear translocation of the transcription factor IRF7. The kinase TBK1 phosphorylates AIP to promote IRF7 binding and the inhibition of the type I interferon (IFN) response. However, it is unknown if AIP expression in innate immune cells is important to suppress type I IFN in the context of RNA virus infection *in vitro* and *in vivo*. In this study we generated myeloid cell-specific AIP conditional knockout mice (*Aip*^fl/fl^xLysM-Cre) to investigate AIP regulation of innate immune signaling in myeloid cells. Bone marrow-derived macrophages (BMDMs) from *Aip*^fl/fl^xLysM-Cre mice had diminished viral replication and increased production of IFNα/b and IL-6 in response to RNA virus infection. AIP-deficient macrophages exhibited increased IRF7 expression and impaired virus-induced IRF7 degradation. AIP interacted with the E3 ubiquitin ligase SOCS1 and enhanced SOCS1 stability and its interaction with IRF7 to promote the proteasomal degradation of IRF7. *Aip*^fl/fl^xLysM-Cre mice exhibited improved survival upon influenza A virus (IAV) infection compared to control mice. Together, these results indicate that myeloid cell-specific AIP suppresses the innate immune response by targeting IRF7.

## Introduction

The aryl hydrocarbon receptor (AhR) interacting protein (AIP; also known as XAP2, ARA9, and FKBP37) functions as a chaperone for AhR to regulate the xenobiotic response^1,2^. AIP and Hsp90 are components of the AhR cytoplasmic complex and together sequester and stabilize AhR in its inactive conformation^3^. AIP also functions as a tumor suppressor in the pituitary gland and loss of function mutations of AIP are associated with pituitary adenomas^4,5^.

Recent studies have revealed new AhR-independent roles of AIP in the regulation of innate and adaptive immunity. Following T-cell receptor activation, AIP binds to the adaptor protein CARMA1 (also known as CARD11) and promotes the formation of the CARMA1-BCL10-MALT1 complex, thus triggering downstream NF-κB signaling and T cell activation^6^. AIP also promotes B-cell proliferation in germinal centers through its interaction with the de-ubiquitinating enzyme, ubiquitin carboxy-terminus hydrolase 1 (UCHL1), which blocks the proteasomal degradation of the transcription factor B-cell lymphoma 6 (BCL6)^7^. Although AIP enhances adaptive immunity through its functions in T and B lymphocytes, it functions as a negative regulator of the innate immune response. AIP binds to the transcription factor IRF7, a master regulator of type I interferon (IFN) production, and prevents its nuclear translocation^8^. The kinase TBK1 phosphorylates AIP on Thr40 to promote IRF7 interaction and inhibition^9^. However, it remains unclear if AIP has functional roles in innate immune cells and *in vivo* in regulating inflammatory responses to virus infection.

Global knockout of AIP in mice leads to embryonic lethality due to cardiac malformations^10^. Conditional knockout floxed AIP mice have been generated and hepatocyte-specific deletion of AIP promotes resistance to dioxin-induced hepatotoxicity^11^. In this study, we have generated myeloid cell (i.e., monocytes, macrophages, granulocytes)-specific AIP knockout mice (*Aip*^fl/fl^xLysM-Cre) and demonstrate that AIP suppresses type I IFN production in macrophages through downregulation of IRF7. Furthermore, we demonstrate that AIP interacts with the E3 ubiquitin ligase suppressor of cytokine signaling 1 (SOCS1) and promotes its stability and interaction with IRF7 to trigger its degradation. Finally, myeloid cell-specific AIP-deficient mice infected with IAV had decreased mortality compared to wild-type (WT) mice.

## Results

### AIP inhibits RNA virus-induced type I IFN and IL-6 in primary macrophages

To understand the functional roles of AIP in innate immunity in myeloid cells we generated myeloid cell-specific AIP knockout mice by crossing *Aip*^fl/fl^ mice with LysM-Cre mice. These mice appeared similar physically to WT mice with no significant differences in weight (Fig. S1). Spleen sizes were also comparable in *Aip*^+/+^xLysM-Cre and *Aip*^fl/fl^xLysM-Cre mice (data not shown). We next examined myeloid cell populations in the spleens of *Aip*^+/+^xLysM-Cre and *Aip*^fl/fl^xLysM-Cre mice by flow cytometry. The percentage of neutrophils was similar, inflammatory monocytes and macrophages were increased, whereas plasmacytoid dendritic cells (pDCs) were decreased in *Aip*^fl/fl^xLysM-Cre mice compared to WT controls (Fig. S2). We previously reported that AIP inhibits type I IFN and IRF7 in response to RNA virus infection^8,9^; however, these studies were mainly performed in mouse embryonic fibroblasts (MEFs). To determine if AIP inhibits type I IFN in primary macrophages, we generated bone marrow-derived macrophages (BMDMs) from *Aip*^+/+^xLysM-Cre and *Aip*^fl/fl^xLysM-Cre mice. Flow cytometric analysis revealed comparable differentiation of *Aip*^+/+^xLysM-Cre (WT) and *Aip*^fl/fl^xLysM-Cre (*Aip*^−/−^) BMDMs (Fig. S3). We next infected WT and *Aip*^−/−^ BMDMs with Sendai virus (SeV), an RNA virus that induces copious amounts of type I IFN. The expression of *Ifnα4*, *Ifnb* and *Il6* mRNAs was upregulated in *Aip^−/−^* BMDMs in response to SeV infection (Fig. 1A). There was also a significant increase in the protein expression of IFNα (all 14 mouse IFNα subtypes) and IL-6 in the supernatant of *Aip^−/−^* BMDMs as determined by ELISA (Fig. 1B). Furthermore, infection with a different RNA virus, carboxy terminal GFP tagged vesicular stomatitis virus (VSV-GFP), also yielded increased expression of *Ifnα4*, *Ifnb* and *Il6* mRNAs mRNAs in *Aip*^−/−^ BMDMs (Fig. 1C). Together, these results indicate that AIP suppresses type I IFN and IL-6 in primary mouse macrophages infected with RNA viruses.

**Figure 1.**
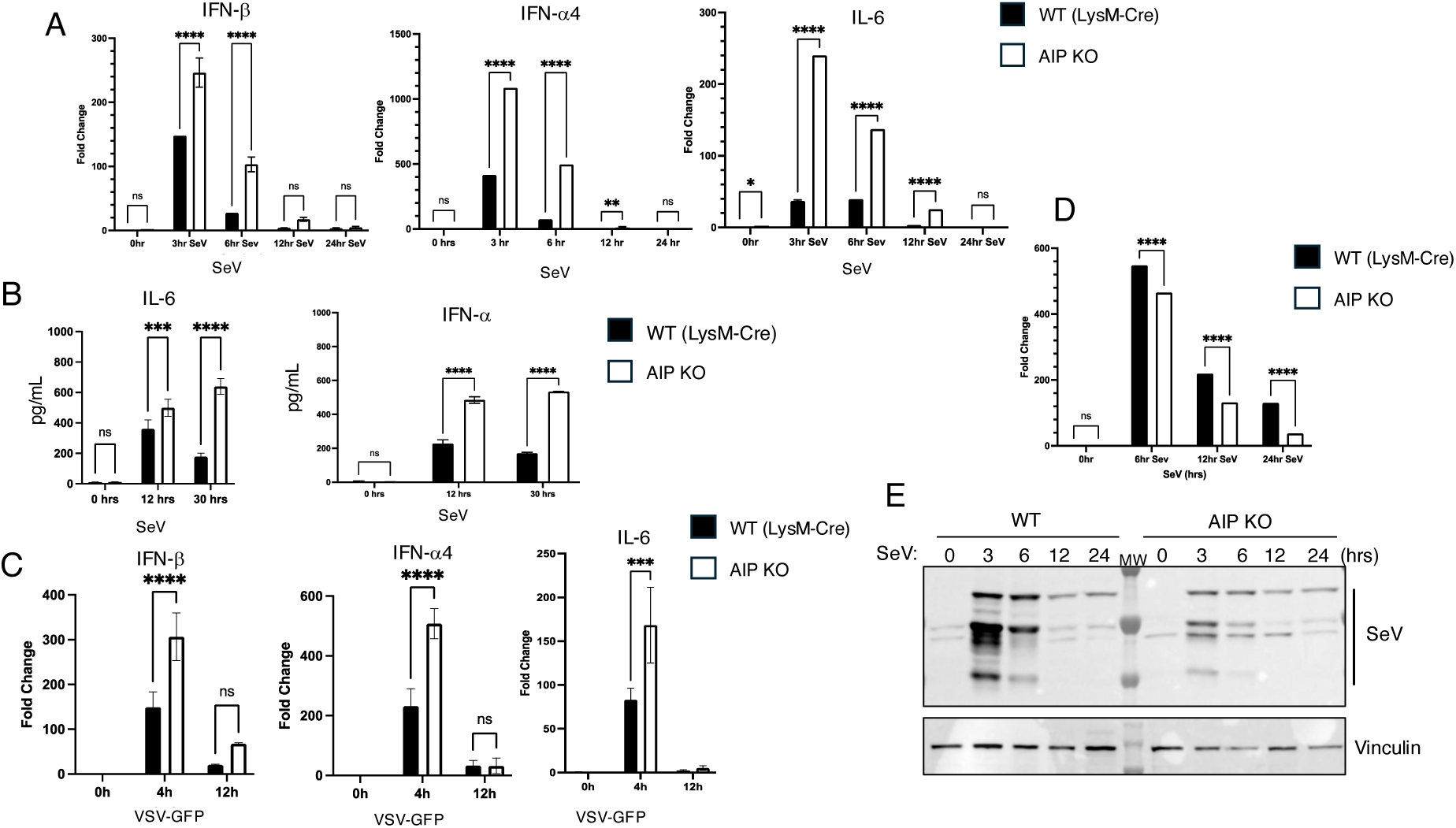
*Aip*^−/−^ BMDMs exhibit increased type I IFN and cytokine production in response to RNA virus infection. **(A)** qRT-PCR analysis of SeV-induced *Ifnb*, *Ifnα4* and *Il6* mRNAs in *Aip^+/+^* and *Aip^−/−^* BMDMs infected with 30 HA/mL SeV for the indicated time points. Fold induction was normalized to uninfected WT BMDMs. **(B)** Supernatants from *Aip^+/+^* and *Aip^−/−^* BMDMs infected with 30 HA/mL SeV for 0, 12, and 30 hrs. were collected and subjected to ELISA for the detection of IL-6 and IFNα (all 14 mouse subtypes). **(C)** qRT-PCR analysis of *Ifnb*, *Ifnα4* and *Il6* mRNAs in *Aip^+/+^* and *Aip^−/−^* BMDMs infected with VSV-GFP (MOI=10) for the indicated time points. Fold induction was normalized to uninfected WT BMDMs. **(D, E)** SeV infection in *Aip^+/+^* and *Aip^−/−^* BMDMs was assessed by qRT-PCR of SeV *P* mRNA expression (D), and western blot analysis of SeV proteins using an anti-SeV polyclonal antibody (E). (Two-way ANOVA, n ≥ 3, * indicates p-value ≤ 0.05, ** indicates p-value ≤ 0.01, *** indicates p-value ≤ 0.001, and **** indicates p-value ≤ 0.0001, ns=not significant).

Type I IFN is critical to restrict viral replication and spread, therefore we hypothesized there would be less viral replication in *Aip*^−/−^ BMDMs. Indeed, the mRNA and protein expression of SeV was significantly diminished in *Aip*^−/−^ BMDMs as measured by qRT-PCR of the *P* gene encoded by SeV and western blotting with a SeV polyclonal antibody (Fig. 1D, E). VSV infection was also diminished in *Aip*^−/−^ BMDMs (data not shown). Therefore, *Aip*^−/−^ BMDMs exhibit resistance to RNA virus infection, likely due to increased expression of type I IFN.

### AIP does not regulate the phosphorylation of RLR signaling proteins but suppresses IRF7 mRNA and protein levels

RNA-virus-induced type I IFN production is mediated by the RIG-I-like receptor (RLR) pathway. The main RLRs are the DExD/H box RNA helicases RIG-I and melanoma differentiation associated gene-5 (MDA5), which recognize 5’-triphosphate dsRNA and long dsRNA, respectively^12^. RIG-I/MDA5 sensing of viral RNAs promotes activation of the mitochondrial adaptor MAVS that recruits TRAF E3 ubiquitin ligases and the kinase TBK1 to activate the transcription factor IRF3^13^. The kinase IKKb and NF-κB transcription factor are also activated downstream of MAVS to induce expression of proinflammatory cytokines, such as IL-6^14^. IRF3 mainly induces type I IFN in the early phase of the innate antiviral immune response whereas IRF7 is an IFN-stimulated gene (ISG) that mediates the later amplification phase of the innate immune response^15^. Although IRF7 was likely the main target of AIP during RLR signaling, we examined upstream signaling events by western blotting with phospho-specific antibodies. Loss of AIP did not affect SeV-induced phosphorylation of TBK1, IRF3, or IRF7 (Fig. 2A, B). Likewise, loss of AIP did not affect SeV-induced phosphorylation of p65/RelA or total p65/RelA protein levels (Fig. 2A, B). However, *Irf7* mRNA expression was significantly increased in *Aip*^−/−^ BMDMs following SeV infection (Fig. 2C). While IRF3 protein levels were not different between *Aip^+/+^* and *Aip*^−/−^ BMDMs infected with SeV (Fig. 2B), there was significantly more IRF7 protein in *Aip*^−/−^ BMDMs after SeV infection (Fig. 2D).

**Figure 2.**
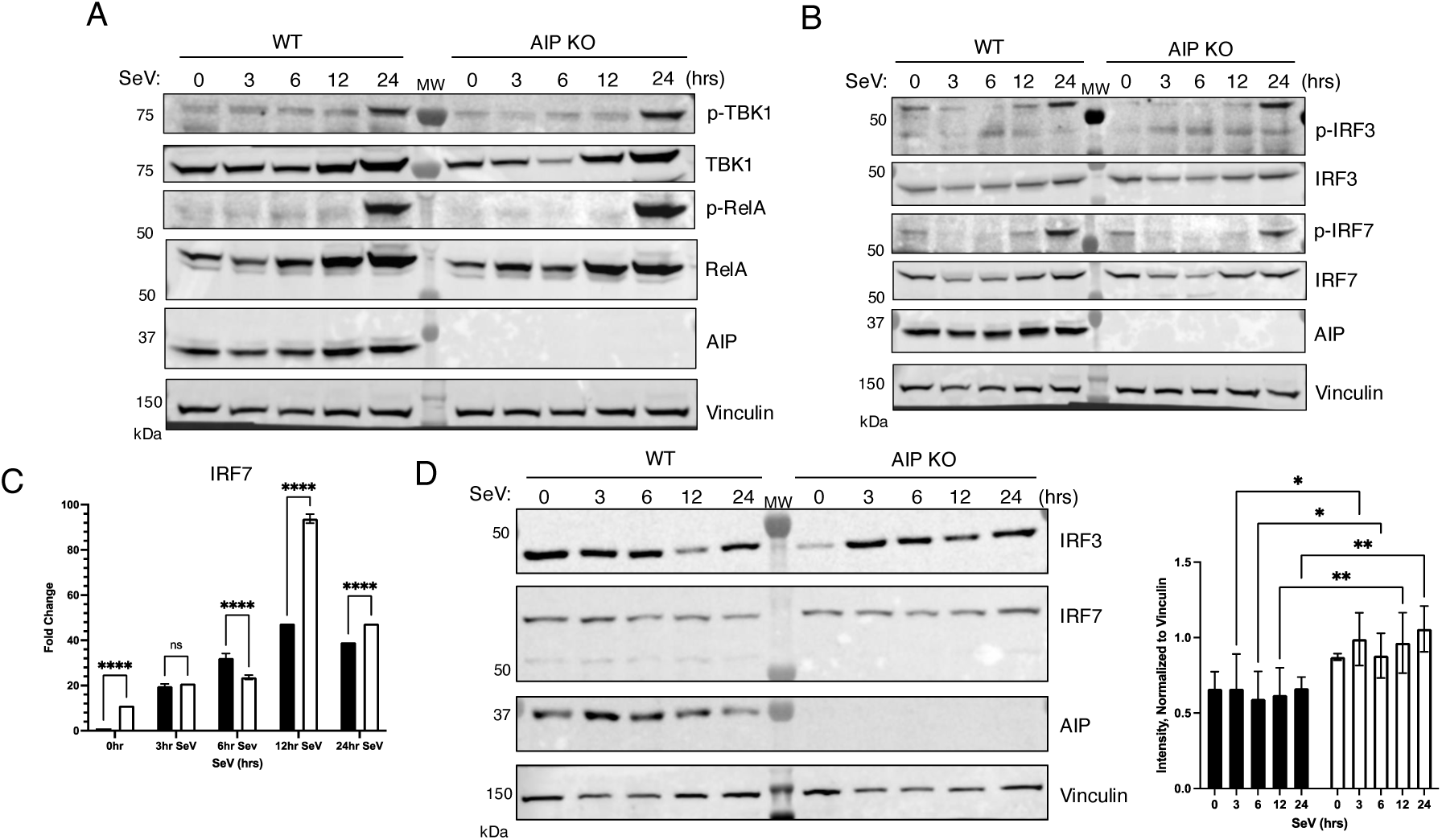
AIP does not regulate the phosphorylation of RLR signaling proteins but inhibits IRF7 expression. **(A,B)** *Aip^+/+^* and *Aip^−/−^* BMDMs were infected with 30 HA/mL SeV for the indicated time points and lysates were subjected to western blotting to examine phosphorylation of the indicated proteins. **(C)** qRT-PCR of *Irf7* mRNA in *Aip^+/+^* and *Aip^−/−^* BMDMs infected with 30 HA/mL SeV for the indicated time points. **(D)** Western blot analysis of IRF3 and IRF7 using lysates from *Aip^+/+^* and *Aip^−/−^* BMDMs infected with 30 HA/mL SeV for the indicated time points. Band intensity was quantified and normalized to vinculin. (Two-way ANOVA, n ≥ 3, * indicated p-value ≤ 0.05, ** indicates p-value ≤ 0.01, and **** indicates p-value ≤ 0.0001, ns=not significant).

### AIP enhances SOCS1-mediated proteasomal degradation of IRF7

SOCS1 is an E3 ubiquitin ligase that inhibits JAK/STAT signaling and is induced by IFN in a negative feedback loop^16^. Mice lacking SOCS1 die between 2 and 3 weeks of age due to liver necrosis, severe lymphopenia and macrophage infiltration and inflammation of several organs due to unchecked IFNγ signaling^17^. In addition to its critical roles in inhibiting type I and II IFN signaling, SOCS1 also inhibits IFNα/b expression by targeting IRF7 for proteasomal degradation^18^. Interestingly, a yeast-2-hybrid screen previously conducted by our lab using a mouse macrophage cDNA library and SOCS1 as bait identified AIP as a potential binding partner (data not shown). Therefore, we explored a potential regulation of SOCS1 by AIP. SeV-induced *Socs1* mRNA expression was significantly increased in *Aip*^−/−^ BMDMs (Fig. 3A), likely due to enhanced type I IFN responses (*Socs1* is an interferon-stimulated gene) in the absence of AIP. SOCS1 protein was not detectable basally but was strongly induced in both WT and *Aip^−/−^* BMDMs upon SeV infection and downregulated by 24 hours after infection (Fig. 3B). AIP has been shown to function as a chaperone for many proteins to promote their stabilization^19^. Indeed, overexpression of AIP stabilized SOCS1 as shown by a CHX chase assay (Fig. 3C). Since SOCS1 was shown to interact with IRF7 and trigger its proteasomal degradation^18^, we next sought to determine if AIP regulated SOCS1-mediated degradation of IRF7. Next, a co-IP assay was performed with lysates of cells transfected with Flag-IRF7, Myc-SOCS1 and AIP and infected with SeV for various time points. IP of Flag-IRF7 pulled down AIP and SOCS1, and these interactions were potentiated by SeV infection (Fig. 3D). These data suggest that AIP may form a complex with SOCS1 and IRF7 in response to SeV infection. A CHX chase assay revealed that overexpression of AIP enhanced SOCS1-mediated degradation of IRF7 (Fig. 3E). IRF7 degradation by SOCS1 and AIP was blocked by the proteasome inhibitor MG-132 but not the V-ATPase proton pump inhibitor bafilomycin-A1 (BafA1) (Fig. 3F). Together, our data suggest that AIP and SOCS1 promote IRF7 degradation by the proteasome.

**Figure 3.**
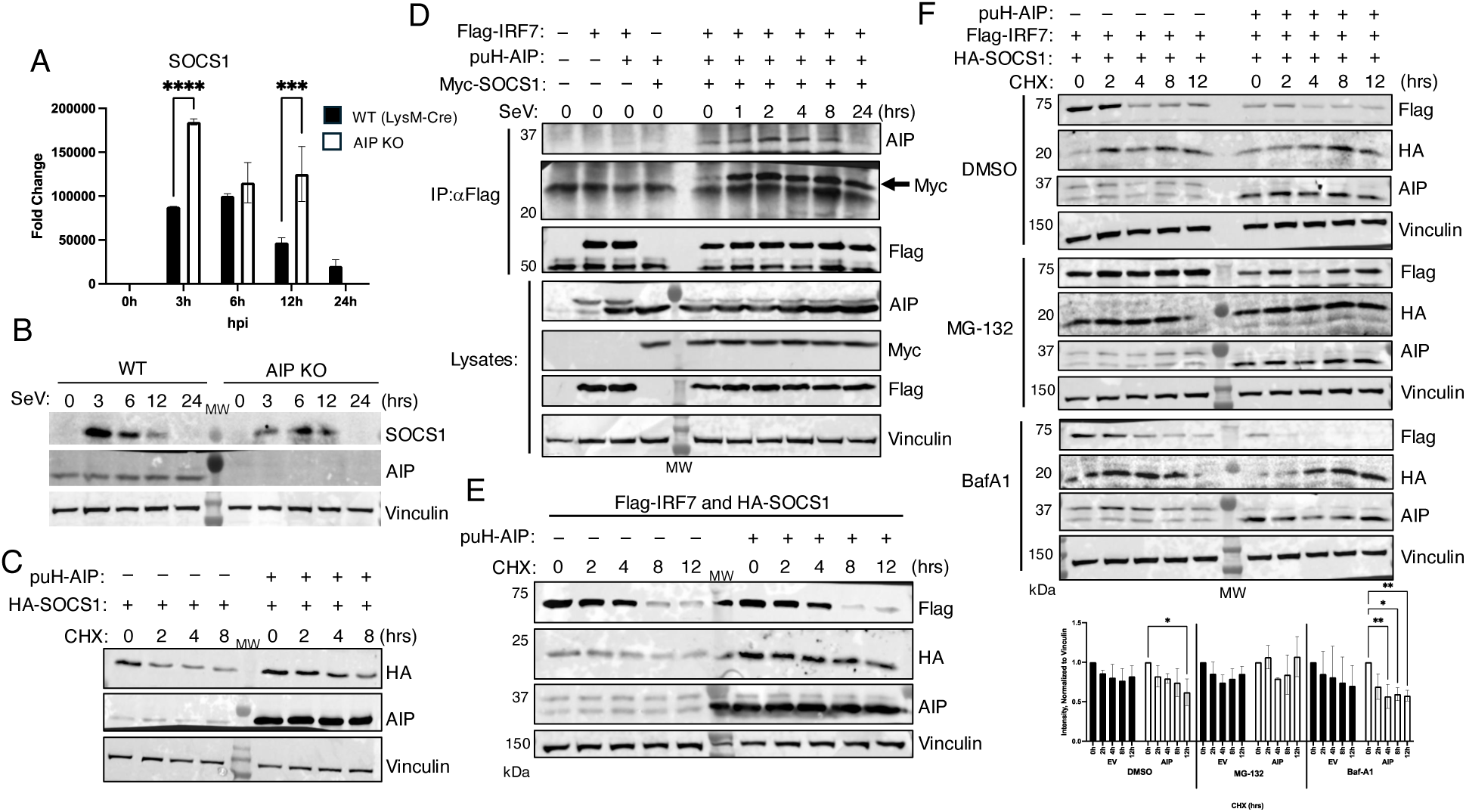
AIP and SOCS1 promote IRF7 proteasomal degradation. **(A)** qRT-PCR of *Socs1* mRNA in *Aip^+/+^* and *Aip^−/−^* BMDMs infected with 30 HA/mL SeV for the indicated time points. Fold induction of *Socs1* was normalized to *Actin* and untreated WT BMDMs. **(B)** Western blot analysis of the indicated proteins using lysates from *Aip^+/+^* and *Aip^−/−^* BMDMs infected with 30 HA/mL SeV for the indicated time points. **(C)** CHX chase assay using lysates from 293T cells transfected with 0.5 μg pUltraHot-AIP and/or 0.25 μg HA-SOCS1 and 24 hrs. later cells were treated with 100 μg/mL cycloheximide for the indicated times. **(D)** co-IP assay and western blotting with the indicated antibodies using lysates from 293T cells transfected with 0.25 μg Myc-SOCS1, 0.25 μg Flag-IRF7, and/or 0.5 μg pUltra-Hot-AIP and infected with 30 HA/mL SeV the following day for the indicated times. **(E)** CHX chase assay using lysates from 293T cells transfected with 0.25 μg HA-SOCS1, 0.25 μg Flag-IRF7, and/or 0.5 μg pUltraHot-AIP and 24 hrs. later cells were treated with 100 μg/mL cycloheximide for the indicated times. **(F)** CHX chase assay using lysates from 293T cells transfected with 0.25 μg HA-SOCS1, 0.25 μg Flag-IRF7, and/or 0.5 μg pUltraHot-AIP and treated with vehicle control (dimethyl sulfoxide), proteasome inhibitor (MG-132, 10 μM), or lysosomal inhibitor (BafA1, 100 nM) together with 100 μg/mL cycloheximide for the indicated times. (Two-way ANOVA, n ≥ 3, * indicates p-value ≤ 0.05, ** indicates p-value ≤ 0.01, *** indicates p-value ≤ 0.001, and **** indicates p-value ≤ 0.0001).

### Myeloid cell-specific AIP-deficient mice have enhanced survival upon IAV infection

Given that AIP-deficient MEFs and BMDMs are resistant to RNA virus infections^8^ (Fig. 1), we next sought to determine if myeloid cell-specific AIP-deficient mice were resistant to IAV. *Aip*^+/+^xLysM-Cre and *Aip*^fl/fl^xLysM-Cre mice were infected intranasally with a sublethal dose of mouse-adapted IAV (PR8). *Aip*^fl/fl^xLysM-Cre mice exhibited significantly decreased IAV-induced mortality (Fig. 4A). Whereas only one of sixteen *Aip*^fl/fl^xLysM-Cre mice died, 50% of *Aip*^+/+^xLysM-Cre died from the same dose of IAV (Fig. 4A). Surprisingly, there were no significant differences in weight loss between *Aip*^+/+^xLysM-Cre and *Aip*^fl/fl^xLysM-Cre mice following IAV infection (Fig. 4B). There was also no difference in IAV NS1 mRNA expression in lung tissues of *Aip*^+/+^xLysM-Cre and *Aip*^fl/fl^xLysM-Cre mice at days 3 and 7 after IAV infection (data not shown).

**Figure 4.**
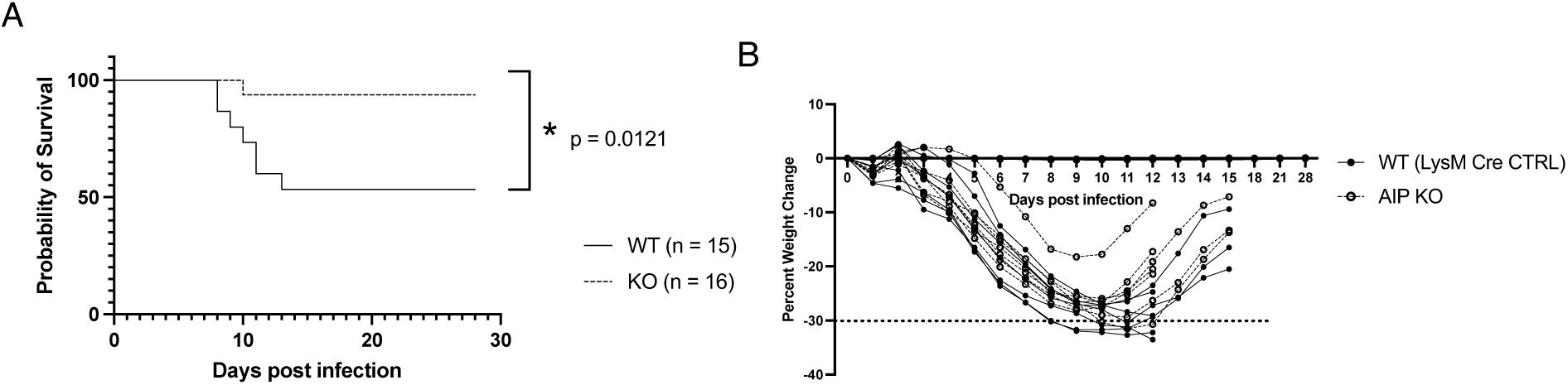
Enhanced survival of myeloid cell-specific AIP-deficient mice infected with IAV. **(A,B)** 12-week-old *Aip*^+/+^xLysM-Cre and *Aip*^fl/fl^xLysM-Cre mice female mice were infected with IAV (2000 FA/mL) and monitored daily for four weeks for survival (A) and weight loss (B).

## Discussion

We previously demonstrated that TBK1 phosphorylates AIP on Thr40 to promote an interaction with IRF7 and prevent its translocation to the nucleus^9^. In this study, we have generated myeloid cell-specific AIP-deficient mice and show that macrophages from these mice have upregulated type I IFN and IL-6 in response to RNA virus infection. We have also identified a novel mechanism of AIP inhibition of IRF7 via SOCS1 whereby AIP interacts with SOCS1 and enhances its stability. AIP may also form a virus-inducible complex with SOCS1 and IRF7 to enhance SOCS1-mediated proteasomal degradation of IRF7. Therefore, AIP utilizes two distinct mechanisms to suppress IRF7: 1) sequestration of IRF7 in the cytoplasm^8,9^, and 2) complex formation with SOCS1/IRF7 to enhance IRF7 degradation.

The SOCS family of proteins function as inducible negative regulators of cytokine signaling^16^. SOCS1 is a critical inhibitor of type I and II IFN signaling by blocking JAK-STAT signaling^20^. SOCS1 has a SOCS box domain that can assemble components of an E3 ubiquitin ligase complex to degrade specific proteins^21^. SOCS1 and SOCS3 can interact with and promote the ubiquitination and proteasomal degradation of IRF7^18^. Our data show that AIP interacts with and stabilizes SOCS1 and may also enhance SOCS1 interaction with IRF7 to trigger its polyubiquitination and degradation by the proteasome. This is analogous to how AIP regulates the anti-apoptotic protein survivin. AIP interacts with and stabilizes survivin^22^, and through its interaction with TOM20 shuttles survivin to the mitochondria to exert its anti-apoptotic function^23^. SOCS1 has also been shown to target RelA/p65 for degradation^24^; however, we did not observe any differences in RelA/p65 phosphorylation or expression in *Aip*^−/−^ BMDMs (Fig. 2). Since we consistently detected increased levels of IL-6 in *Aip*^−/−^ BMDMs infected with SeV (Fig. 1), there is likely enhanced NF-κB activation in the absence of AIP. Future studies should examine IKK phosphorylation/activation, IκBα phosphorylation and degradation and RelA/p65 nuclear translocation in AIP-deficient macrophages in response to SeV infection as well as TNF and IL-1b stimulation. These experiments will reveal the specificity and potential mechanism by which AIP inhibits NF-κB activation. It is plausible that AIP may prevent the nuclear translocation of RelA/p65, similar to AIP-mediated cytoplasmic retention of IRF7.

AIP has been extensively studied as a chaperone protein for AhR and its regulation of AhR stability and signaling in response to dioxin^2^. However, we did not observe any differences in AhR protein levels between *Aip^+/+^* and *Aip^−/−^* BMDMs upon SeV infection (Fig. S4). In addition to metabolizing dioxins, AhR can function as a negative regulator of inflammation and immunity^25^, but there were no differences in AhR activity in *Aip*^−/−^ BMDMs, as measured by the AhR target gene *CYP1B1* expression, in response to SeV infection (Fig. S4). We speculate that in response to virus infection, AIP phosphorylation by TBK1 switches AIP function from AhR signaling to the negative regulation of type I IFN and inflammation by binding to SOCS1 and IRF7. It will be interesting to examine if AIP phosphorylation regulates SOCS1 binding, and therefore in TBK1 knockout cells, SOCS1 targeting of IRF7 may be impaired.

Global *Aip^−/−^* mice die in utero from cardiac malformations^10^. To study the functional roles of AIP in myeloid cells *in vivo* in response to virus infection, we generated myeloid cell-specific AIP-deficient mice. Our *in vivo* studies show that myeloid cell-specific AIP-deficient mice exhibit decreased mortality in response to IAV infection (Fig. 4). However, there were no significant differences in weight loss or IAV NS1 expression in lung tissues in *Aip*^+/+^xLysM-Cre and *Aip*^fl/fl^xLysM-Cre mice (Fig. 4 and data not shown). Therefore, the mechanisms underlying the enhanced survival of IAV-infected *Aip*^fl/fl^xLysM-Cre mice are unclear but could potentially be explained by differences in lung repair, inflammation and/or adaptive immune responses. There is precedence for such a phenotype as mice lacking the absence in melanoma 2 (AIM2) inflammasome infected with IAV had improved survival compared to WT mice but with no differences in viral loads or weight loss^26^. Since *Aip*^−/−^ BMDMs express more IL-6 in response to RNA virus infection, increased IL-6 expression by myeloid cells *in vivo* could potentially promote the survival of IAV-infected *Aip*^fl/fl^xLysM-Cre mice. Previous studies have shown that IL-6 was crucial for lung repair after IAV infection^27^ and mediated host defense against IAV by enhancing neutrophil survival in the lung^28^ and eliciting B cell differentiation and protective antibody responses^29^. Future studies should examine whether increased IL-6 expression accounts for the enhanced survival of *Aip*^fl/fl^xLysM-Cre mice infected with IAV.

In summary, our study has identified a novel mechanism of AIP regulation of IRF7 and the type I IFN response in macrophages, in which AIP stabilizes SOCS1 and promotes the proteasomal degradation of IRF7 to suppress type I IFN production. In addition, myeloid cell-specific AIP-deficient mice were resistant to IAV infection.

## Materials and Methods

### Cells, plasmids, and reagents

HEK293T cells were obtained from ATCC (CRL-3216). HA-SOCS, Myc-SOCS1 and pUltra-Hot-AIP were previously described^9,30^. Flag-IRF7 was provided by Dr. John Hiscott (Istituto Pasteur Italia). CHX, BafA1 and MG-132 were purchased from Millipore-Sigma. pUltra-Hot was a gift from Malcolm Moore (Addgene plasmid # 24130; http://n2t.net/addgene:24130; RRID:Addgene_24130)

### Mice

All animal studies were performed within an American Association for the Accreditation of Laboratory Animal Care-certified barrier facility at Pennsylvania State College of Medicine. All animal work was approved by the Institutional Animal Care and Use Committee (Protocol No 01844). Mice were housed under specific pathogen-free conditions and were provided with food and sterile water ad-libitum and maintained on a 12-hour light/12-hour dark cycle.

LysM-Cre (strain #004781) and *Aip*^fl/fl^ mice (strain #013195) were obtained from Jackson Laboratories. Myeloid cell-specific *Aip* knockout mice were generated by crossing *Aip*^fl/fl^ mice with LysM-Cre mice. Mouse genotyping was performed by obtaining genomic DNA from ear punches using Extracta DNA prep (QuantBio). PCR was carried out using RedExtract-N-Amp PCR Ready Mix (Millipore-Sigma) with the following primers: *Aip*^fl/fl^ primers: forward 5’- CAATCCCCCACTGTCACTTT -3’ and reverse 5’- TCACCCCTCCCACTGACTAC -3’ and Cre: forward 5’- GCCTGCATTACCGGTCGATGCAACGA -3’ and reverse 5’- GTGGCAGATGGCGCGGCAACACCATT -3’.

### Cell culture and transfections

HEK293T cells were cultured in Dulbecco’s modified fetal bovine serum supplemented with 10% heat-inactivated fetal bovine serum and 1% penicillin-streptomycin. Mouse BMDMs were isolated from the femur and tibia of 8-to 12-week-old LysM-Cre and *Aip*^fl/fl^xLysM-Cre mice and differentiated into macrophages using L929 conditioned complete media (50% DMEM, 30% L929 media, 20% heat inactivated FBS, and 1% penicillin-streptomycin) over four days. Cells were incubated at 37°C and 5% CO_2_. DNA transfections of HEK293T cells were performed using GenJet Plus (SignaGen) according to the manufacturer’s instructions.

### Co-immunoprecipitation assays and western blotting

Cells were lysed with RIPA buffer (50 mM Tris pH 7.4, 2% SDS, 5% glycerol) containing HALT protease and phosphatase inhibitor cocktail (ThermoFisher Scientific). Protein concentration was estimated using the BCA protein assay kit (ThermoFisher Scientific). For co-immunoprecipitation (co-IP) assays, 400 μg of protein lysate was incubated with mouse anti-Flag or mouse anti-Myc antibody (Proteintech) (1:200). The next day, lysates were incubated with 20 μL of protein A/G PLUS-agarose beads (Santa Cruz Biotechnology) for 4 hrs. Beads were washed, and lysates were dissociated from beads according to the manufacturer’s instructions. Lysates were prepared in 4x NuPAGE LDS sample buffer with NuPAGE reducing agent (ThermoFisher Scientific) and heated at 75°C for 5 min. Lysates were run on 6% to 15% gradient SDS-PAGE gels and transferred onto nitrocellulose membranes using the Trans-Blot Turbo Transfer System (Bio-Rad Laboratories). Antibodies used for western blotting are listed in Table S1. Western blots were developed with SuperSignal West Pico PLUS Chemiluminescent Substrate (ThermoFisher Scientific) and imaged with an Azure 600 Imager. Band intensity was quantified using ImageJ.

### Quantitative real-time PCR

BMDMs were seeded in a six-well plate at 10^6^ cells/well. Total RNA was isolated using the Zymo Quick RNA kit (Zymo Research). cDNA was synthesized using M-MLV reverse transcriptase (Invitrogen) according to the manufacturer’s instructions. Quantitative real-time PCR reactions were performed using 5x PowerUp SYBR Green Master Mix (Applied Biosystems). Primers are listed in Table S2. All reactions were run in triplicate on QuantStudio 3 Real-Time PCR System (ThermoFisher Scientific) and analyzed via Design and Analysis software (Applied Biosystems). Fold induction (bbCt) was calculated by normalizing threshold cycle values of genes of interest to b-actin and then relative to untreated control (*Aip*^+/+^), as previously described^9^.

## ELISA

Supernatants were collected and centrifuged at 500xg for 5 min. Mouse IFNα and IL-6 ELISA kits were purchased from PBL Assay Science and Biotechne (R&D systems), respectively. ELISAs were performed according to the manufacturers’ instructions.

### Virus infections

Sendai virus (SeV) (Cantell strain) was purchased from Charles River Laboratories. VSV-GFP was provided by Dr. Glen Barber (Ohio State University). IAV strain PR8 was provided by Dr. Zissis Chroneos (Penn State University).

For *in vitro* virus infections, cells were serum starved for 1 hr., then incubated in serum-free media containing SeV (30 HA/mL) or VSV-GFP (MOI = 10) for 1 hr. with rocking every 15 min. Cells were replenished with complete medium following incubation with virus-containing medium.

For *in vivo* IAV infections, 12-week-old female mice were anesthetized with an intraperitoneal injection of ketamine (100 mg/kg) and xylazine (10 mg/kg). Mice were infected intranasally with 40 μL of 2000 fluorescent focus units (FFC) of the mouse adapted H1N1 strain A/Puerto Rico/8/13 (PR8). Infected mice were weighed and monitored daily for clinical symptoms as detailed previously^31^.

### Flow cytometry

Differentiation of BMDMs was assessed by flow cytometry using > 10^6^ cells. Cells were incubated in Fc Receptor Blocking Solution (TruStain FcX Plus; Biolegend) for 10 min. and then incubated for 20 min. in a mixture of the following: 7-AAD viability dye (1:300), mouse F4/80 (1:150), and mouse CD11b (1:300) (Table S3), which were diluted in Flow Cytometry Staining Buffer (ThermoFisher Scientific). Cells were then washed three times with the staining buffer and analyzed by flow cytometry using a 23-color BD FACSymphony.

Myeloid cell populations were assessed from spleens of 8-week-old LysM-Cre and *Aip*^fl/fl^xLysM-Cre mice. Spleens were homogenized by pressing the tissue through a cell strainer and washing with PBS. The cells were then pelleted and resuspended in 5 mL of 1x RBC lysis buffer (BioLegend) and incubated for 5 min. Cells were washed with PBS before flow cytometric analysis. Mouse antibodies are listed in Table S3. Compensation and fluorescent minus one (FMOs) were calculated. Flow cytometry analysis was carried out using FlowJo software.

### Statistical analysis

Statistical analysis was performed using GraphPad Prism 9.3.1, Error bars represent standard deviation of multiple samples. The specific statistical test used for experiments is indicated in the Figure Legends and statistical significance is indicated as * p ≤ 0.05, ** p ≤ 0.01, *** p-≤ 0.001, and **** p-≤ 0.0001.

## Acknowledgements

We thank Dr. Glen Barber for VSV-GFP. We thank John Tawil for experimental contributions to this study. We acknowledge the Penn State College of Medicine Flow Cytometry Core and the Pulmonary Immunology and Physiology Core. The publication of this study was supported by the National Institute of Allergy and Infectious Diseases: NIH R21AI153731. The Flow Cytometry Core (RRID:SCR_021134) services and instruments used in this project were funded, in part, by the Pennsylvania State University College of Medicine via the Office of the Vice Dean of Research and Graduate Students and the Pennsylvania Department of Health using Tobacco Settlement Funds (CURE). The content is solely the responsibility of the authors and does not necessarily represent the official views of the University or College of Medicine. The Pennsylvania Department of Health specifically disclaims responsibility for any analyses, interpretations, or conclusions.

---

**Figure S1.**
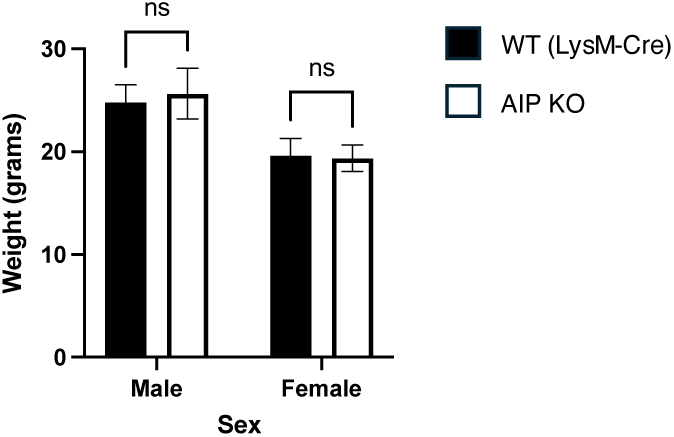
Myeloid-cell specific AIP-deficient mice have similar weights compared to LysM-Cre control mice. Weights of 12-week-old male (n=12) and female (n=24) *Aip*^+/+^xLysM-Cre and *Aip*^fl/fl^xLysM-Cre mice. There was no significant difference between genotypes. ns=not significant.

**Figure S2.**
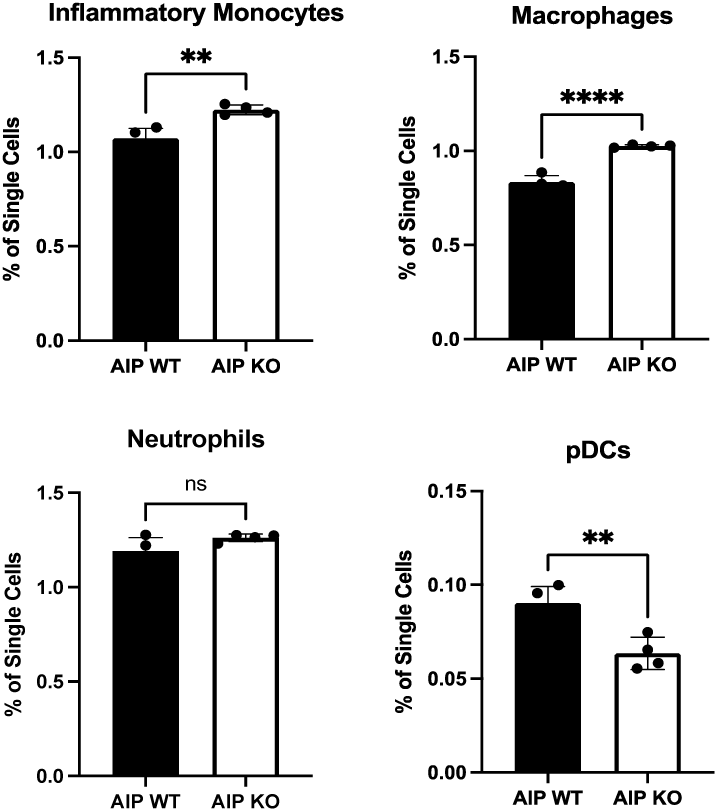
Inflammatory cell populations in spleens from LysM-Cre and *Aip*^fl/fl^xLysM-Cre mice. Flow cytometric analysis of myeloid cell populations in spleens from LysM-Cre and *Aip*^fl/fl^xLysM-Cre mice. Inflammatory monocytes are CD11b^hi^CD11c^−^MHCII^−^Ly6C^hi^Ly6G^−^ F4/80^+^. Macrophages are CD11b^hi^CD11c^−^MHCII^−^ CD8^-^Ly6C^lo^Ly6G^-^F4/80^+^. Neutrophils are CD11b^hi^CD11c^−^MHCII^−^CD8^−^Ly6C^+^Ly6G^+^F4/80^−^. pDCs are CD11b^−^ CD11c^+^MHCII^lo^Ly6C^hi^Ly6G^−^. (n=4, Unpaired Student’s *t*-test, ns=not significant, ** indicates p < 0.01, and **** indicates p < 0.0001).

**Figure S3.**
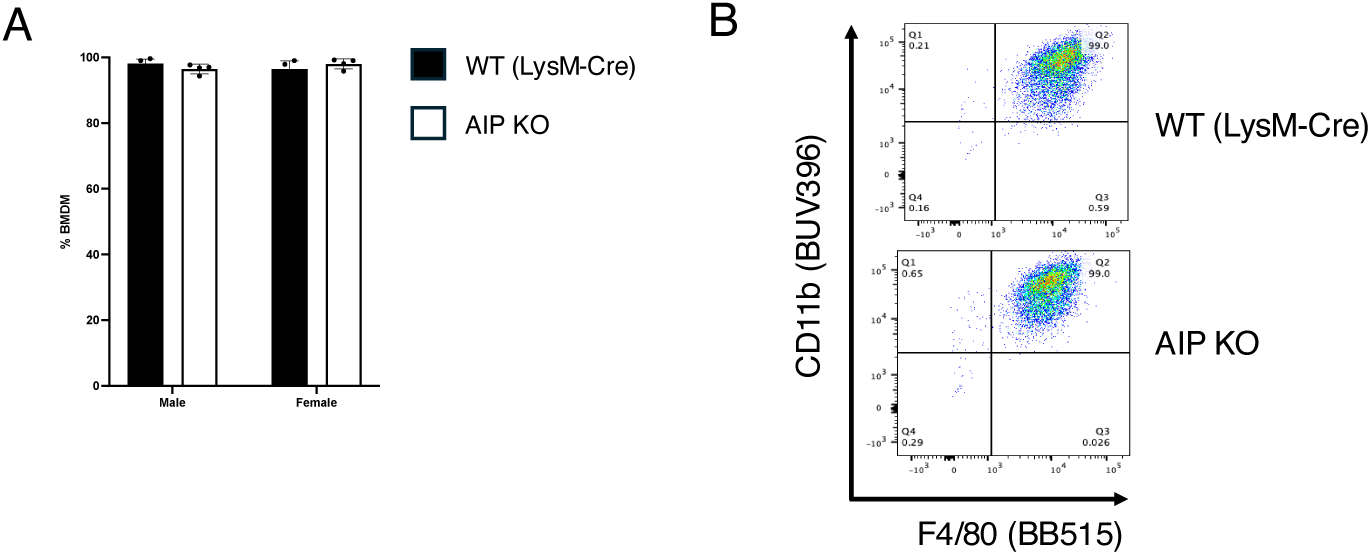
Comparable differentiation of WT and *Aip^−/−^* BMDMs. **(A)** The differentiation of BMDMs was assessed by flow cytometric analysis of CD11b and F4/80. **(B)** A representative flow cytometry grid is shown. There was no significant difference in CD11b^+^F4/80^+^ expression between *Aip^+/+^* and *Aip^−/−^* BMDMs.

**Figure S4.**
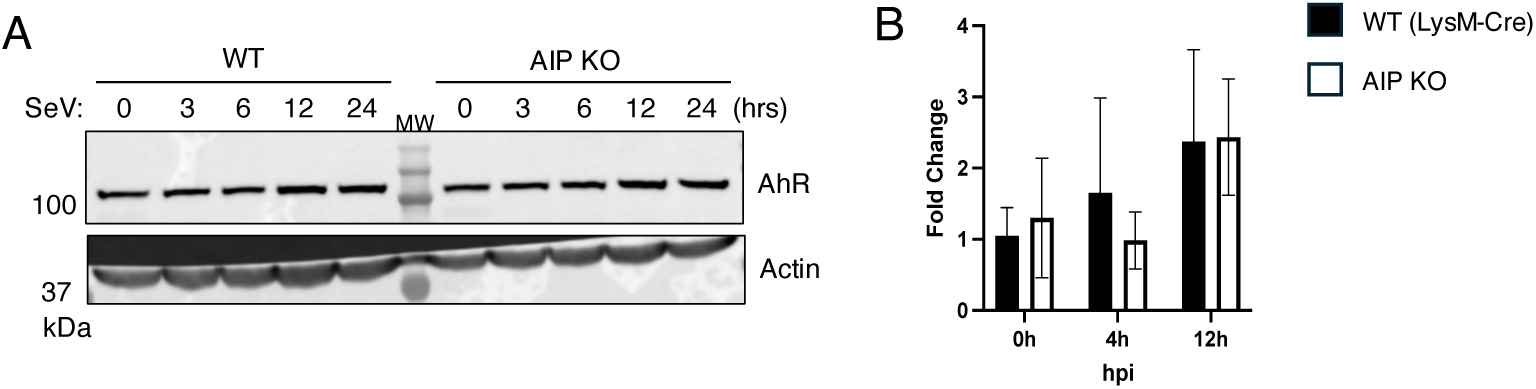
AhR expression and activation in *Aip*^−/−^ BMDMs infected with SeV. **(A)** *Aip^+/+^* and *Aip^−/−^* BMDMs were infected with 30 HA/mL SeV for the indicated times. Lysates were subjected to western blotting for the indicated proteins using lysates from *Aip^+/+^* and *Aip^−/−^* BMDMs. **(B)** qRT-PCR of the AhR target gene *Cyp1b1* in *Aip*^+/+^ and *Aip*^−/−^ BMDMs following SeV infection.

**Table S1.** Antibodies for Western Blotting.

| **Table S1. Antibodies for Western Blotting** |  |  |
| --- | --- | --- |
| **Antibody** | **Catalog Number** | **Company** |
| Mouse anti-Actin | sc47778 | Santa Cruz |
| Mouse anti-AIP | sc59730 | Santa Cruz |
| Mouse anti-Flag | 66008-4-Ig | Proteintech |
| Mouse anti-HA | 3724S | Cell Signaling Technology |
| Mouse anti-Myc | 60003-2-Ig | Proteintech |
| Mouse anti-Vinculin | sc73614 | Santa Cruz |
| Rabbit anti-SOCS1 | 55313S | Cell Signaling Technology |
| Rabbit anti-phospho-TBK1 | 5483S | Cell Signaling Technology |
| Rabbit anti-TBK1 | 3806S | Cell Signaling Technology |
| Rabbit anti-phospho-IRF3 | 4947S | Cell Signaling Technology |
| Rabbit anti-IRF3 | 4302S | Cell Signaling Technology |
| Rabbit anti-phospho-IRF7 | 5184S | Cell Signaling Technology |
| Rabbit anti-IRF7 | AHP1180 | BioRad |
| Rabbit anti-phospho-p65 | 3033S | Cell Signaling Technology |
| Rabbit anti-p65 | 6956S | Cell Signaling Technology |
| Rabbit anti-phospho-STAT1 | 9167S | Cell Signaling Technologies |
| Rabbit anti-STAT1 | 14994T | Cell Signaling Technology |
| Rabbit Anti-AhR | 28727-1-AP | Proteintech |
| ECL Anti-mouse IgG (Secondary) | NA931V | Cytiva |
| ECL Anti-mouse IgG (Secondary) | NA934V | Cytiva |

**Table S2.**
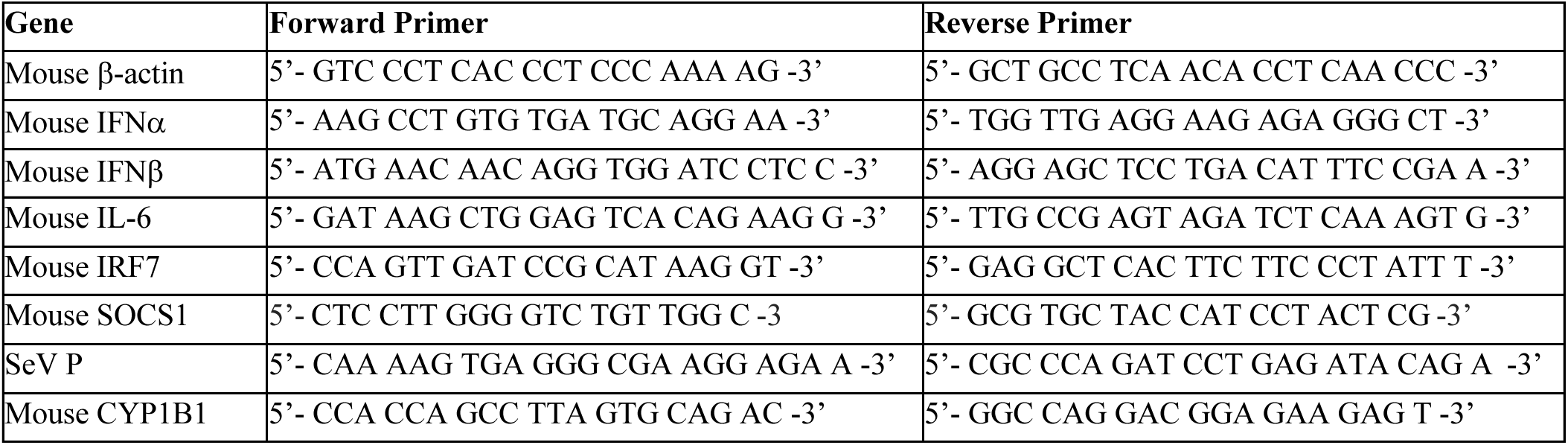
Primers for qRT-PCR.

**Table S3.** Antibodies for flow cytometry.

| **Table S3. Antibodies for flow cytometry** |  |  |  |  |
| --- | --- | --- | --- | --- |
| **Antibody** | **Color** | **Clone** | **Catalog Number** | **Company** |
| 7-AAD | PE Cy5 |  | 420403 | BioLegend |
| F4/80 | Alexa 488 | Clone BM8 | 123120 | BioLegend |
| CD11b | BUV395 | Clone M1/70 | 563553 | BioLegend |
| CD4 | Alexa 488 | GK15 | 100423 | BioLegend |
| CD11c | PE | N418 | 117308 | BioLegend |
| CD19 | BV750 | 6D5 | 115561 | BioLegend |
| CD80 | BV711 | 16-10A1 | 104743 | BioLegend |
| CD86 | BV421 | PO3 | 105123 | BioLegend |
| Ly6C | APC | HK14 | 128015 | BioLegend |
| Ly6G | BV605 | 1A8 | 127639 | BioLegend |
| MHC-II | APC/Fire 750 | AF6-120.1 | 116423 | BioLegend |
| SiglecF | BUV737 | IRNM44N | 367-1702-82 | BioLegend |

## Notes

### Competing Interest Statement

The authors have declared no competing interest.

